# Humans predictively process high-level information during active natural vision

**DOI:** 10.64898/2026.11.27.769638

**Authors:** Luca Kämmer, Carmen Amme, Philip Sulewski, Martin N. Hebart, Tim Kietzmann

## Abstract

Prediction is considered a fundamental principle of perception. However, evidence for predictive visual processing comes largely from paradigms that establish explicit predictive relationships, leaving unclear whether the brain spontaneously generates feature-level predictions during natural viewing. We analyzed magnetoencephalography data recorded concurrently with eye movements as participants freely explored thousands of natural scenes. Using linear deconvolution to disentangle overlapping neural responses, we demonstrate that high-level conceptual information derived from artificial neural networks is reflected in neural activity approximately 50–100 ms before fixation onset, whereas low-level perceptual information emerges only after fixation onset. Source analyses localized these predictive high-level signals to higher-order cortical regions. Consistent with feature-level prediction, neural responses 400-600 ms after fixation onset varied with how predictable the high-level features of fixation targets were from preceding fixations. Together, these findings suggest that during active natural vision, the brain continuously predicts high-level content of upcoming visual input, shaping subsequent processing.

## Introduction

Visual perception is influenced by prior expectations about our visual environment. Studies using controlled sequences of stimuli have shown that expectations about upcoming visual input can influence our perception and associated neural responses [1–7]. However, testing predictive processing of upcoming visual input in natural settings is challenging. Overlapping neural responses and shared visual features make it difficult to distinguish anticipation of upcoming fixation content from ongoing processing of currently visible or previously fixated information. Therefore, much of the evidence for predictive visual processing comes from highly constrained paradigms. In these paradigms, participants are exposed to a small number of stimuli that are made predictable through repeated exposure or learned cue-target associations and analyses focus mostly on the violations of predictions instead of prediction itself [2, 8–11]. As these conditions differ from natural visual experience, it remains unclear whether feature-level predictions arise during free viewing of natural scenes without experimentally imposed predictive relationships.

Natural vision offers opportunities to anticipate upcoming visual input. During natural vision, visual information is actively sampled through sequences of fixations [12–16] and upcoming fixation targets are embedded in spatial and semantic context [17]. Additionally, fixation selection itself is guided by high-level content, with observers preferentially sampling semantically relevant regions [12, 18–23]. This context-guided selective sampling may allow the brain to anticipate the content of an upcoming fixation. Yet much of our understanding of visual processing is derived from random sequences of static images presented while participants maintain stable fixation [24–26]. While such paradigms facilitate experimental control and the isolation of specific visual features, they cannot directly address predictions associated with natural eye movements. It also remains unclear whether predictions about upcoming fixation content concern low-level visual properties, high-level content, or both.

To determine the role of predictions of upcoming information during natural eye movements, we analyzed a large-scale dataset in which five participants explored 4080 natural scenes while eye movements and MEG signals were recorded concurrently[27]. We show that high-level image features were associated with neural activity before fixation onset, while low-level features only showed association after fixation onset. Furthermore, these predictive modulations were localized to higher-level cortical regions. To assess whether these effects reflect feature-level predictions, we tested whether post-fixation neural activity tracked the predictability of the fixated content. Post-fixation neural responses varied with the predictability of the fixated content, providing converging evidence for feature-level predictions across saccades during natural vision.

## Results

### Feature dimensions are decodable over time

To test whether neural activity before fixation onset reflects the visual features of the upcoming fixation target, we needed to characterize the visual features at each fixation.

To this end, we extracted image crops centered on each fixation location (Fig. 1a) to focus the analysis on the central visual field. We then described the visual content of these crops using a 49-dimensional embedding of interpretable dimensions derived from similarity judgments (SPoSE). These dimensions capture both low-level and highlevel visual features based on human similarity judgments [28]. SPoSE dimensions have been shown to account for a significant portion of explainable variance in neural responses to images in both fMRI [29] and MEG [30], making them well-suited to track how visual features are encoded across fixations. To predict the dimensions for each fixation crop we used the same approach as previous papers. Dimensions are estimated based on CLIP-ResNet embeddings [31, 32] (Fig. 1b). Fig. 1c illustrates the SPoSE dimensions extracted from fixation crops within an example scene.

**Fig. 1.**
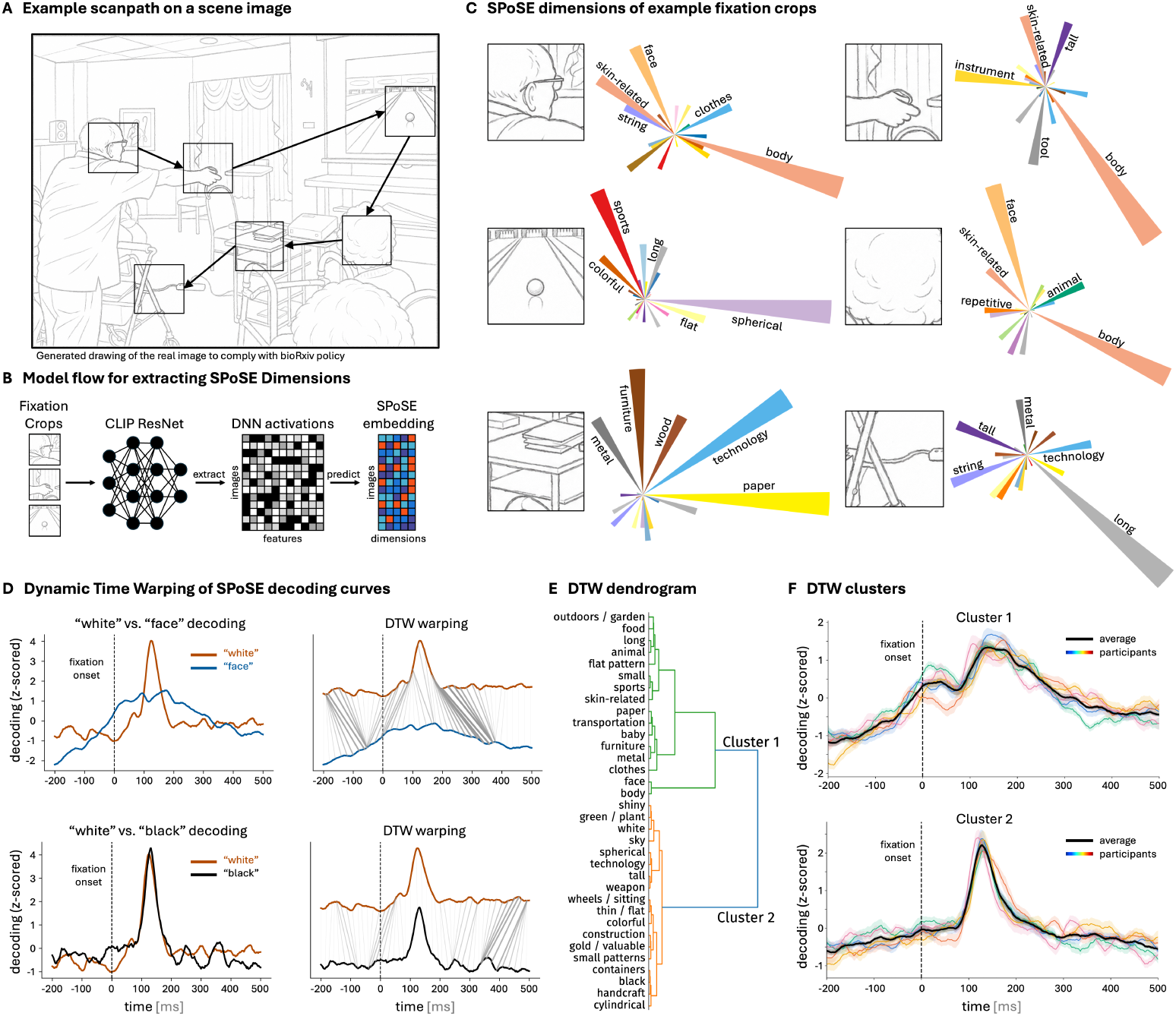
Decoding SPoSE dimensions from continuous MEG data during free viewing. **A)**, Example scanpath on a scene image. **B)**, Extraction of SPoSE dimensions from fixation crops using the CLIP model. **C)**, Predicted SPoSE dimensions extracted from fixation crops. **D)**, Shows the DTW approach for two pairs of decoding time curves. SPoSE dimensions were decoded from the MEG signal around fixation onset. The left side shows z-scored curves on top of each other, the right side shows the warping cost for one dimension to resemble the other. **E)**, Shows the dendrogram, which results from taking the warping costs as a distance measure. **F)**, Depicts the z-scored decoding curves averaged across participants. The colored lines represent the individual participants, while the black lines show the average across them. The shaded lines are the standard error across dimensions. Illustrates that some dimensions show an increase in decoding accuracy before fixation onset (Cluster 1), while others show increased decoding only well after fixation onset (Cluster 2), consistently across all 5 subjects. All images in this figure were substituted with generated drawings to comply with bioRxiv policy.

To test whether neural effects differed between high- and low-level visual features, we grouped the dimensions into low-level / perceptual (color-, shape-, and texture-related) and high-level / conceptual dimensions (categorical, animacy-, and human-related), similar to Kramer et al. (2023)[33]. This grouping was based on the labels that were assigned to each dimension by human ratings (for all the labels, see Hebart et al. (2020)[28]).

To assess whether neural signatures of these dimensions can be observed around fixation onset, we fitted multiple linear regression models to estimate the association between MEG sensor activation patterns and each dimension at each time point. This generated decoding timeseries from −200 to 500 ms around fixation onset, reflecting how well the MEG signal at each time point predicts the feature content of the fixation crop. Most dimensions showed characteristic decoding curves, replicating previous work done on passive viewing data [30] (Supplementary Fig. 1). Decoding accuracy peaked around 100 ms after fixation onset for most dimensions, but time courses differed markedly between dimensions: some showed elevated decoding before fixation onset, while others only increased afterwards. To distinguish these patterns, we applied dynamic time warping [30] as the basis for a clustering approach (Fig. 1d), excluding dimensions with peak decoding below r = 0.05. To determine the optimal number of clusters, we computed the silhouette score, which indicated a two-cluster solution as the best fit (silhouette = 0.38). The two cluster time courses can be seen in Fig. 1f. Cluster 2 showed a sharp increase peaking at 128 ms (SD = 7 ms) after fixation onset followed by a rapid decline. Cluster 1 showed an earlier rise beginning before fixation onset and peaking at 144 ms (SD = 19 ms), with a more gradual decline afterwards. This pattern was consistent across all participants (Fig. 1f). Notably, Cluster 1 contained a higher proportion of high-level dimensions, while Cluster 2 contained more low-level dimensions, with 68% of dimensions following this pattern (Fisher’s exact test, p = 0.043, odds ratio = 4.4) (Fig. 1e). This suggests that high-level features of upcoming fixation crops may influence neural processing before fixation onset, while low-level features are only encoded after the eyes land.

### Linear deconvolution shows predictive high-level feature modulation

These results suggest that the brain may anticipate high-level features of upcoming fixation targets. However, two limitations of the decoding approach complicate this interpretation. First, the SPoSE dimensions are not mutually orthogonal across all fixation crops. For example, the “face” dimension is negatively correlated with the “long” (r = −0.28, SD = 0.001) and the “flat pattern” dimensions (r = −0.18, SD = 0.006), which may lead to an overlap of the decoding patterns of these dimensions (Supplementary Fig. 1). Second, since the decoding model is fitted independently for each time point, it cannot distinguish the neural response to the current fixation from residual processing of the previous one. Since consecutive fixation crops are likely to share features because of the shared spatial statistics within a given scene, pre-fixation decoding could reflect carry-over from the preceding fixation rather than genuine prediction of the upcoming one.

To address these limitations, we modeled the continuous MEG signal as the linear sum of overlapping event-related responses across all fixations (Fig. 2a) using the Unfold toolbox [34]. By time-expanding the design matrix to include predictors for each time point around each fixation event (−300 to +800 ms), the model simultaneously estimates the neural response at each time point while aiming to control for overlap with preceding and following fixations. The model’s intercept captures the average MEG response time-locked to fixation onset, while the beta estimates for each SPoSE dimension reflect how that dimension modulates the neural signal at each time point and sensor, independently of all other dimensions and of previous fixations. To further control for oculomotor confounds, we additionally included saccade amplitude, saccade velocity, and pre and post-saccadic fixation position as regressors. The model was run separately for each MEG magnetometer sensor in the dataset.

**Fig. 2.**
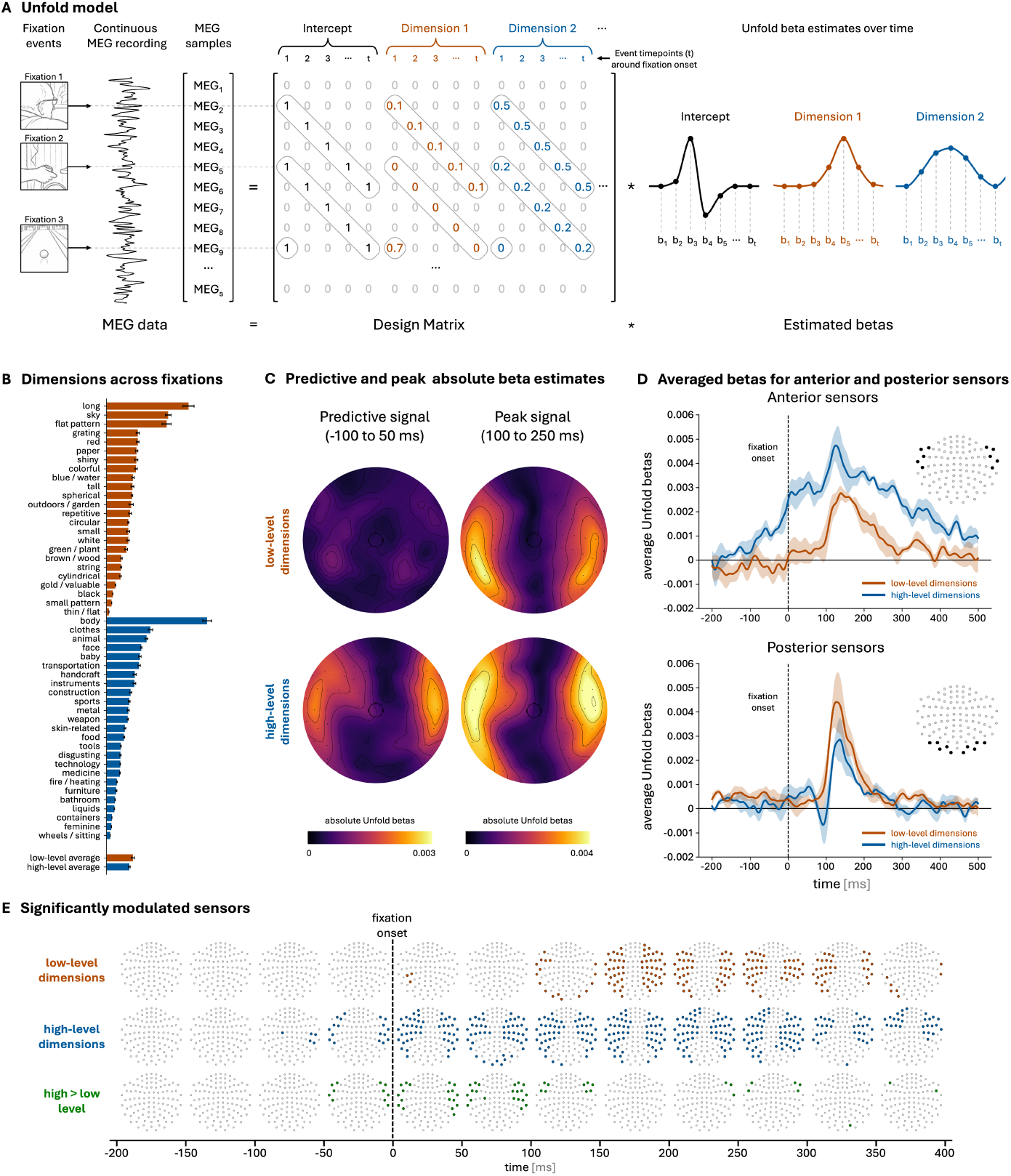
Temporal signal deconvolution reveals high-level dimensions predictively modulating the MEG signal. **A)**, Conceptual representation of how Unfold generates beta estimates of each dimension using deconvolution to isolate the overlapping effects of each fixation event. **B)**, Shows how prevalent each dimension is, by averaging the dimension’s loading across all fixation patches of a given subject. Error bars represent the standard error across the 5 subjects. **C)**, Shows the magnitude of modulation of low- and high-level dimensions at all sensors averaged across the predictive (−100 to 50 ms) and the peak window (100 to 250 ms) **D)**, Shows the Unfold betas averaged across 10 representative posterior and anterior sensors. Time courses were aligned before averaging to avoid cancellation. Shaded areas indicate the standard error of the mean. **E)**, Sensors that showed significant modulation for low- or high-level dimensions and sensors that were modulated significantly more by high-compared to low-level dimensions. The betas were binned to 50 ms and significance was computed using a linear mixed model across the 50 sessions (10 per participant). The figure shows sensors that deviated significantly from 0 with a threshold of p *<* 0.05 after Benjamini-Hochberg multiple comparison correction across sensors and time bins. All images in this figure were substituted with generated drawings to comply with bioRxiv policy.

To assess how low- and high-level features modulate the neural signal around fixation onset, we averaged the beta estimates across the low-level and high-level dimensions for each sensor and time point (Supplementary Fig. 2). Critically, even after controlling for overlap between consecutive fixations, the same pattern of results emerged as in the decoding analysis. High-level dimensions modulated neural responses before fixation onset while low-level dimensions only did so afterwards (Fig. 2c). We tested this by binning the Unfold betas into bins of 50 ms and testing if they significantly diverged from 0 at each sensor (*p <* 0.05, FDR-corrected across sensors and time bins; Fig. 2e). Low-level dimensions showed sustained significant modulation after 100 ms, while high-level dimensions first modulated sensors between −100 and −50 ms. Furthermore, high-level dimensions modulated neural activity significantly stronger compared to low-level dimensions between −50 and 150 ms (*p <* 0.05, FDR-corrected across sensors and time bins; Fig. 2e). Additionally, while both low- and high-level dimensions showed modulation of occipito-temporal sensors after 100 ms, predictive high-level modulation was only present in more anterior, fronto-temporal sensors (Fig. 2e). This difference is illustrated by beta time courses averaged across representative sensors in each region (Fig. 2d).

To assess whether these results generalize beyond the SPoSE feature space and the specific low/high-level dimension split we used, we repeated the Unfold analysis using features extracted from AlexNet [35], in which early and late layers have been shown to capture low- and high-level visual features, respectively [36–38]. We extracted 25 principal components from an early layer (pooling layer 1) and 25 from a late layer (pooling layer 3) using PCA to obtain a number of dimensions comparable to SPoSE. Similarly to our main analysis, late-layer features were associated with neural activity at anterior sensors before fixation onset, while early-layer features were associated with neural activity only after fixation onset (Supplementary Fig. 3). These results confirm the predictive high-level effect and show that this generalizes from the behaviorally relevant SPoSE dimensions to image-derived AlexNet features. As an additional control, we repeated the analysis using two low-level image properties computed directly from the fixation crops, luminance and edge density. Neither feature showed significant pre-fixation modulation, further supporting the absence of predictive modulation for low-level visual features (Supplementary Fig. 4).

### Predictive high-level signals localize to higher-order cortex

Pre-fixation associations with high-level features were concentrated at anterior sensors. This pattern suggested a possible contribution from higher-order cortical regions. To examine the cortical sources of these associations, we applied the same linear deconvolution analysis to beamformer source-reconstructed MEG data, estimating beta time courses across 8,196 cortical vertices. As before, we averaged beta estimates across low- and high-level SPoSE dimensions, respectively, and examined the magnitude of feature-related modulation in two time windows: a predictive window (−100 to 50 ms) and a peak window (100 to 250 ms) (Fig. 3a). We defined the predictive window based on prior work showing that feedforward visual responses are detectable no earlier than 50-100 ms after stimulus onset [39–41], meaning that modulation occurring before 50 ms after fixation onset is unlikely to be caused by feedforward processing of the current fixation and rather reflects anticipatory activity.

**Fig. 3.**
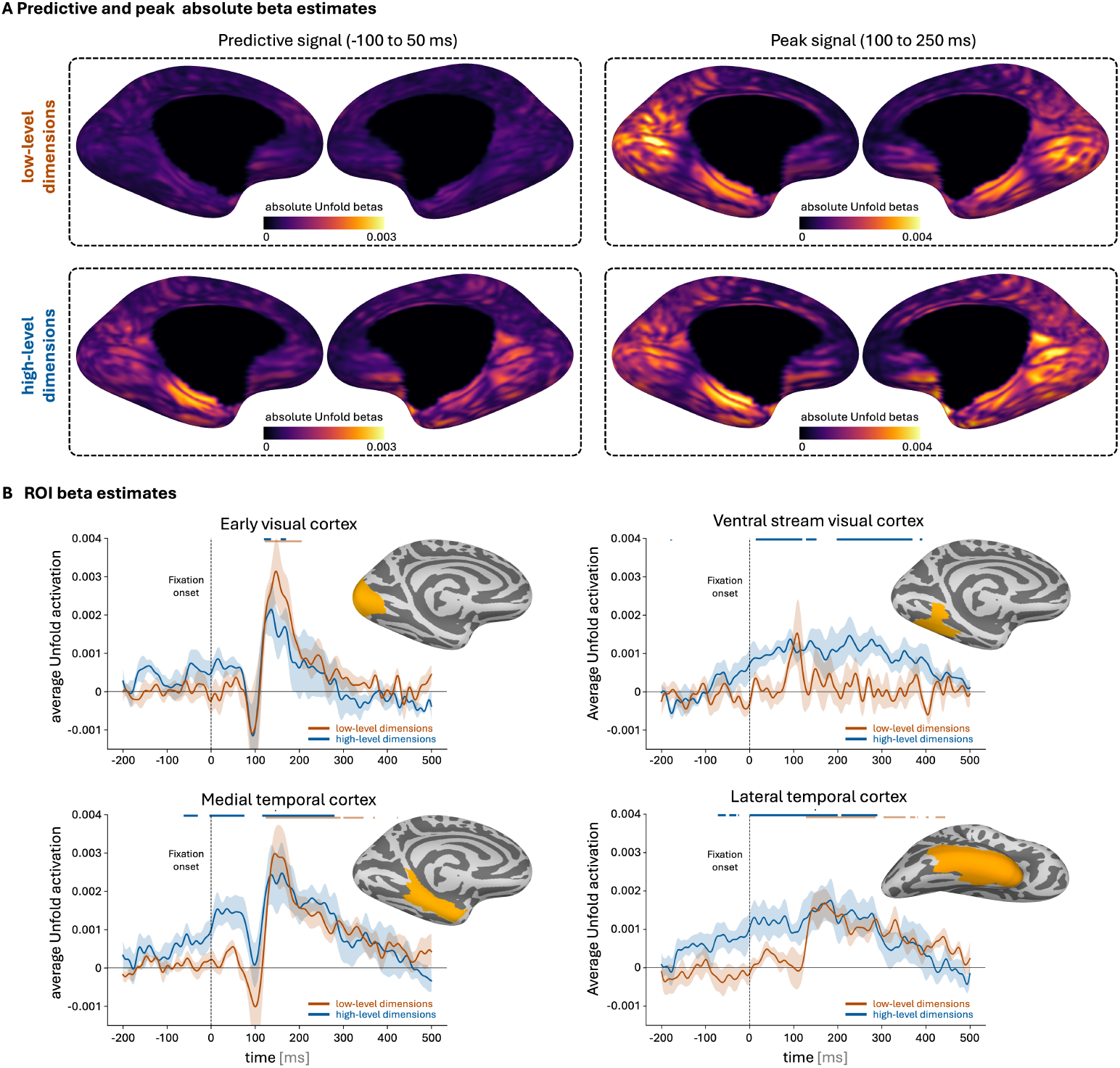
Predictive high-level information in higher, not lower visual cortex. **A)**, Modulation of low- and high-level dimensions from a medial view, averaged across the predictive (−100 to 50 ms) and the peak window (100 to 250 ms). **B)**, Beta activity extracted from relevant areas along the visual hierarchy. Significance was computed using a linear mixed model across the 50 sessions (10 per participant) and FDR-corrected (Benjamini-Hochberg) across all time points. Time points where the beta value differed significantly from 0 (p *<* 0.05) are indicated with a line above the graph. Shaded areas indicate the standard error across all 5 participants.

As expected, the source-level results showed no significant modulation of low-level dimensions during the predictive window, while high-level dimensions showed modulation in higher-order visual and medial temporal regions (Fig. 3a). To characterize this pattern more precisely, we examined the Unfold beta time courses averaged across regions of interest along the visual hierarchy (Fig. 3b). Early visual cortex showed modulation only after approximately 100 ms post-fixation onset for both dimension groups. Ventral regions showed somewhat earlier activation for high-level dimensions, shortly after fixation onset. The earliest signal was found in medial temporal cortex, which showed pre-fixation modulation for high-level dimensions beginning around 80 ms before fixation onset (*p <* 0.05, FDR-corrected across time points). These results indicate that the anticipatory high-level signal is present in later cortical areas rather than early visual cortex, consistent with a predictive mechanism [42–45].

### Less predictable fixation crops elicit modulated post-fixation neural responses

The previous analyses showed that high-level features of upcoming fixation targets were associated with neural activity before fixation onset, with this anticipatory signal localized primarily to higher-order regions. What remains open is whether this predictive information is also related to visual processing after fixation onset, as expected under predictive processing. If the anticipatory high-level signal identified before fixation is functionally relevant to subsequent visual processing, then neural responses after fixation should depend on how predictable the high-level features of the fixation target were from the preceding visual context. Since we observed pre-fixation modulation specific to high-level features, we expected a predictability-dependent response to occur at occipital sensors between 400 and 600 ms after fixation onset, which reflects a time window that has reliably been associated with sensitivity to high-level or semantic predictability in both vision and language research [46–53].

To obtain an estimate of how predictable each fixation target was from the data, we used the Glimpse Prediction Network (GPN) [54]. GPN is a neural network model trained self-supervised to predict the high-level visual features of a fixation crop from the integrated features of all previous fixation crops in a scene. Specifically, the model extracts the feature embedding of each fixation crop using ResNet50 and feeds these into a recurrent LSTM architecture to predict the embedding of the next fixation crop (Fig. 4a). The model also receives the relative saccade displacement vector as input, inspired by efference copies in the brain [55–57]. The cosine distance between the model’s predicted embedding and the true embedding of each fixation crop provides an estimate of how unexpected each fixation target is given the preceding scene context. We refer to this quantity as the GPN prediction error (Fig. 4b,c). If the brain generates high-level feature predictions before each saccade, fixation targets that are harder to predict, indexed by high GPN prediction error, should therefore be associated with modulated neural responses between 400 and 600 ms after fixation onset.

**Fig. 4.**
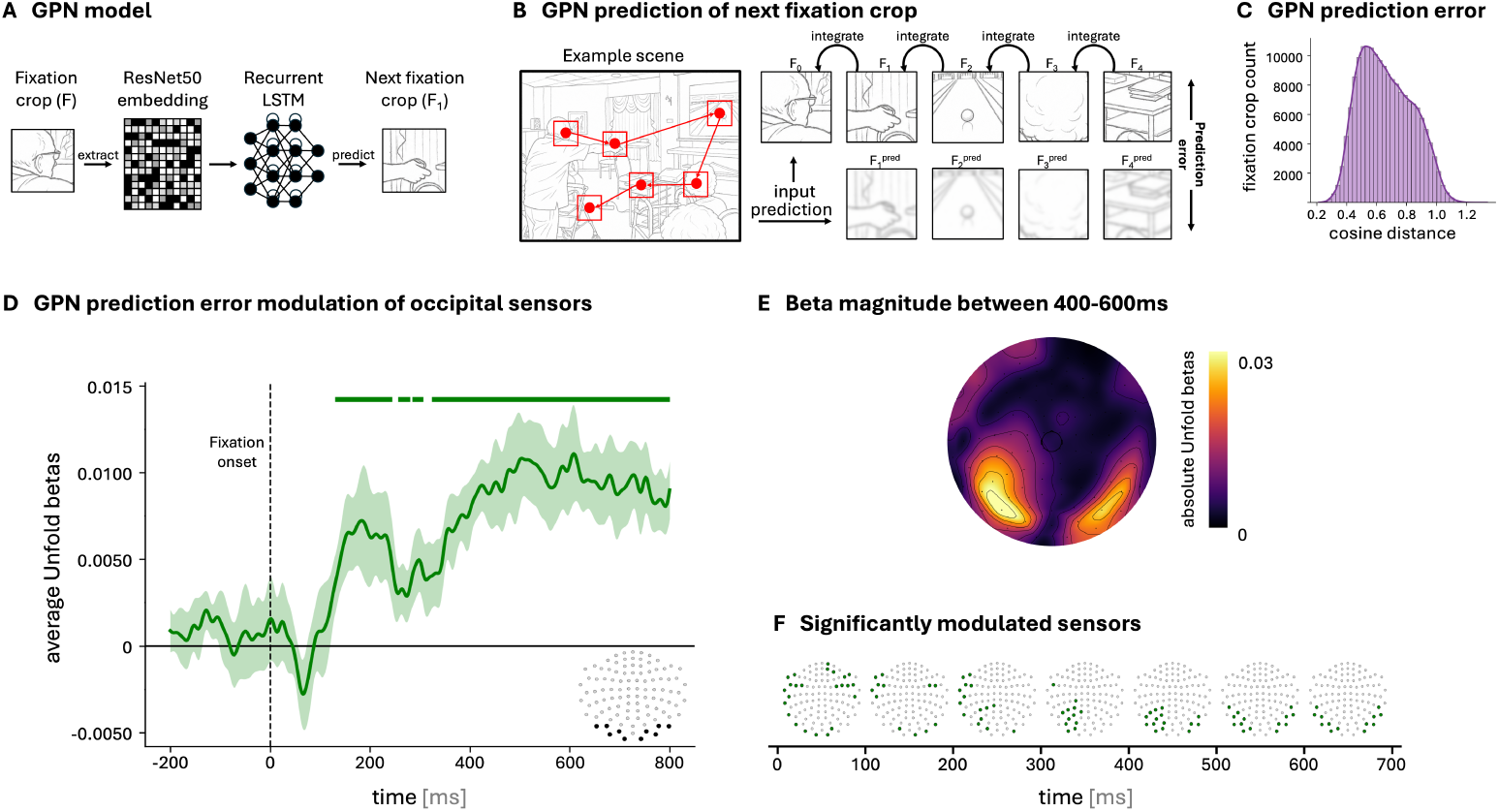
Predictability of high-level features is associated with post-fixation neural responses. **A)**, Shows the conceptual model architecture of GPN. **B)** Illustrates how GPN prediction error is computed, by predicting the embedding of a fixation crop and comparing that prediction to the true embedding of that fixation crop. Shows a conceptual representation of a predicted crop, not a real prediction. **C)**, Distribution of GPN prediction error across all fixation crops. **D)**, Shows Unfold betas estimating how GPN modulates neural activation around fixation onset, averaged across the same 10 posterior sensors used previously **E)**, Shows the absolute modulation of GPN prediction error averaged across 400 to 600 ms after fixation onset. **F)**, Depicts the sensors and time bins at which GPN prediction error significantly modulates Unfold betas. Just as previously the betas were binned to 50 ms and significance was computed using a linear mixed model across the 50 sessions (10 per participant). The figure shows sensors that deviated significantly from 0 with a threshold of p *<* 0.05 after Benjamini-Hochberg multiple comparison correction across sensors and time bins. All images in this figure were substituted with generated drawings to comply with bioRxiv policy.

As this hypothesis concerned the timing and sensor-level distribution of the postfixation predictability effect, we used the sensor-space MEG data for this analysis. To test this hypothesis, we estimated the effect of GPN prediction error on the MEG signal using the same Unfold framework as before. To isolate this effect from potential oculomotor and visual confounds, we included a comprehensive set of control regressors, comprising saccade amplitude, saccade velocity, pre- and post-saccadic fixation position, luminance and edge density of the fixation crop, and all SPoSE dimensions. These regressors control for measured properties of the fixation crops and eye movements that may covary with GPN prediction error. Additionally, we included the similarity between consecutive fixations, computed by correlating their SPoSE dimensions, to control for neural adaptation effects [9, 58, 59].

The results confirmed our hypothesis. Occipital sensors showed significant bilateral modulation by GPN prediction error approximately 400 ms after fixation onset (*p <* 0.05, FDR-corrected across sensors and time bins; Fig. 4f). The signal peaks around 600 ms after fixation onset. This result indicates that anticipatory high-level information may actively contribute to subsequent visual processing after fixation onset. Together with the pre-fixation high-level modulation observed in the preceding analyses, this provides converging evidence for predictive processing of high-level visual content across saccades during natural free viewing.

## Discussion

Using a large-scale dataset of joint MEG and eye-movement recordings, the present study demonstrates that neural activity, recorded during viewing of natural stimuli, is shaped by the high-level content of upcoming fixations. In particular, we found that high-level visual features modulated neural responses from approximately 50-100 ms before fixation onset, while low-level features only did so after fixation onset. This temporal dissociation was accompanied by a spatial dissociation: pre-fixation modulation was localized to anterior and medial temporal regions rather than early visual cortex. Closing the link to a predictive role of this anticipatory activity, post-fixation neural responses varied with how predictable the high-level content of each fixation was from preceding gaze history. Together, these findings suggest that humans predict high-level rather than low-level features of saccade targets during natural viewing.

The most parsimonious alternative explanation for pre-fixation neural modulation is that upcoming saccade targets are visible in peripheral vision before the eyes move, and that this peripheral input, rather than active prediction, drives the effects we observe. Although peripheral preview can affect neural processing in free viewing [60–64], several aspects of our findings argue against a purely peripheral preview-based interpretation. First, the pre-fixation modulation was specific to high-level features, while low-level features showed no pre-fixation modulation. Peripheral preview would be expected to first drive low-level, then high-level features [61], following the typical temporal hierarchy of visual processing [65]. Second, the pre-fixation signal was localized to anterior and medial temporal regions rather than early visual areas, which would be the primary recipients of peripheral visual input [66]. Third, fixation crops that were less predictable, based on previous fixation crops, produced modulated predictability-dependent neural responses 400-600 ms after fixation onset, consistent with a high-level prediction error [46–53]. Such a response indicates that the brain formed a high-level prediction about the fixation content. Relatedly, previous work using linear deconvolution during natural free viewing has shown that category-specific neural effects can emerge before fixation onset and depend on fixation history [67]. This convergent pattern of evidence supports the interpretation that the pre-fixation effects we observe reflect predictive processing of the upcoming fixation content rather than peripheral processing alone. This prediction is likely based on fixation history and additionally informed by general scene context available to the participants.

A key finding of the present study is that predictive modulation before fixation onset was selective for high-level, conceptual features while low-level visual features (color, texture, and shape) only modulated neural responses after fixation onset. This dissociation was consistent across both the SPoSE (Fig. 2) and AlexNet (Supplementary Fig. 3) analyses, and when running the analysis on luminance and edge density, directly computed from the fixation crops (Supplementary Fig. 4). It is consistent with previous reports of high-level specificity in predictive neural signals [11, 42, 68–70] and is consistent with the interpretation that predictions in the brain are conceptual, rather than detailed sensory templates at the pixel-level [42, 71].

However, these findings cannot fully rule out the existence of low-level feature prediction. While we did not find evidence for the processing of low-level features before fixation onset in our analyses using SPoSE, AlexNet, or luminance and edge-density features, it has been reported during controlled experiments [7, 72–74]. The reason may be that, during natural viewing of scene images, the pre-saccadic fixation location contains meaningful visual information rather than a blank fixation cross, which may reduce anticipatory low-level processing. Furthermore, low-level visual features are encoded in a retinotopically precise manner, and anticipatory representations of such features may be difficult to detect before foveal input is available. What we can conclude is that the brain predicts high-level visual features even under the complex, unconstrained conditions of natural free viewing.

The analysis of source-level data revealed that pre-fixation high-level modulation was localized to high-level cortical regions, particularly medial temporal lobe (MTL) rather than early visual cortex. These findings are consistent with a growing body of evidence that implicates MTL in high-level visual processing [75–78]. In particular, perirhinal cortex, which is considered the apex of the ventral visual pathway and part of MTL [79], has been shown to code for object-specific semantic information [80, 81] and complex visual object discrimination [82]. Furthermore, regions in the MTL have been associated with predictive visual processing specifically [44, 45, 83], including predictive processes during free viewing [84]. Our finding that high-level predictive information is present in medial temporal regions is therefore consistent with a role of the MTL in predictive visual processing.

While conventional MEG recordings have been shown to have lower sensitivity to deeper brain structures (such as MTL), less superficial brain activity can be detected with MEG under appropriate conditions [85, 86]. To that end we used beamformer source reconstruction, which is less biased towards superficial sources compared to other methods. Furthermore, we estimated beta coefficients from approximately 200,000 fixation events using linear deconvolution. By accounting for temporal overlap between event-related responses, linear deconvolution may help reveal weaker contributions from deeper brain structures that would otherwise be obscured by overlapping activity. Lastly, during the MEG recording head stabilization was improved by using individually fitted foam cases, thereby minimizing head movement artifacts and improving source localization accuracy [87]. While the strongest predictive high-level modulation was localized to MTL, the spatial limitations of MEG source reconstruction prevent us from attributing this activity unequivocally to the MTL itself. Nevertheless, the localization to higher-order cortical regions is consistent with our interpretation that these signals reflect high-level predictive processing, and is also in line with previous findings implicating the MTL in visual prediction.

The number of participants used in this study is limited, similar to other deep sampling datasets [24, 25, 88], which may limit generalizability. However, without deep sampling, the present results could likely not have been found, because linear deconvolution requires a large number of samples. Still, future work should extend this to larger samples to show that the results can generalize. Furthermore, while we find converging evidence for high-level feature prediction across saccades, future work should use direct manipulation to further investigate this effect. Additionally, our estimate of predictability of a fixation crop is purely model-based, using GPN. Finally, MEG source reconstruction in medial temporal regions has inherent spatial limitations, and combining MEG with fMRI or invasive recording in future studies would provide more precise localization of the deep sources implicated here.

Taken together, the present findings show the value of investigating visual perception in a more natural setting, allowing us to assess predictive processes across eye movements. This study suggests that under naturalistic vision the brain actively predicts high-level information to anticipate upcoming input before it arrives.

## Methods

### Data collection and pre-processing

#### Participants

MEG data were collected from five right-handed participants (three female; age range, 26–33 years; mean age, 27.8 years). All participants were native German speakers, reported no history of neurological or psychiatric illness, and had normal or corrected-to-normal vision. Written informed consent was obtained at the beginning of every experimental session, as well as on the days of scanning and fitting of the individualized head stabilizers. The study was approved by the ethics committee of the University Medical Center Leipzig and was conducted in accordance with the Declaration of Helsinki. Participants received financial compensation for their participation.

#### Stimuli

The stimulus set comprised 4,080 images selected from the Natural Scenes Dataset (NSD) [24], which contains 73,000 natural scene images from the Microsoft Common Objects in Context (MS COCO) dataset [89]. Images were pseudo-randomly sampled from NSD under the constraint that semantic content was represented as evenly as possible. K-means clustering was applied to all NSD scene captions, resulting in 60 semantic clusters, from which equal numbers of images were drawn. Of the 4,080 selected images, 25% (n = 1,020) were assigned to caption trials, in which participants produced a spoken description following image presentation. Caption images were selected according to two additional constraints. First, caption images included the 100 images defined by Allen et al. (2022) [24] as maximally spanning semantic space (“special100”). Second, images from the NSD “shared1000” set, which are presented to all participants, were included whenever doing so did not violate the balanced cluster distribution. All participants viewed the same set of caption and non-caption images, but image order was randomized separately for each participant.

#### Experimental Procedure

Each participant completed 10 recording sessions distributed across 2–5 weeks. During the experiment, participants freely viewed images, and on 25% of trials they were additionally instructed to produce a spoken caption after image presentation. Caption and non-caption images were presented in random order. Participants were seated at a viewing distance of 70 cm from the display. Stimuli subtended 28.54° × 21.61° of visual angle. At the beginning of the first session, participants received instructions in German and were shown 10 example images, each accompanied by five example captions. At the start of every session, the eye tracker was calibrated using a standard 9-point procedure in which participants fixated dots presented sequentially at nine screen locations. Each trial began with a centrally presented white fixation cross on a gray background for at least 1 second. This fixation period also served as a driftcorrection step for fine eye-tracker calibration. Trials were restarted whenever gaze deviated by more than 1.3° from the fixation cross. Each image was presented for 4 seconds, during which participants were allowed to inspect it freely. On 75% of trials, the subsequent fixation cross immediately marked the beginning of the next trial. On the remaining 25% of trials, image presentation was followed by a centrally presented black microphone symbol on a gray background for 1 second, indicating that the participant should verbally describe the image. Audio recording began when the microphone cue disappeared and continued for 8 seconds, after which the next trial commenced. Stimuli were presented in blocks of 30 trials. The first session consisted of 10 blocks (300 trials total), whereas each of the remaining nine sessions consisted of 14 blocks (420 trials per session). After every 3–4 blocks, the eye tracker was recalibrated and participants were given a short break. All spoken captions were manually transcribed after completion of the experiment.

#### Eye-Tracking Data Acquisition and Preprocessing

Eye movements were recorded at 500 Hz using an EyeLink 1000 system (SR Research; https://www.sr-research.com). Raw eye-tracking signals were processed using the saccade detection algorithm described by Engbert and Mergenthaler (2006)[90]. Velocity vectors were computed as sample-to-sample differences separately for the horizontal and vertical gaze coordinates. These x- and y-velocity components were normalized by their respective standard deviations, and speed was defined as the norm of the resulting sample-wise velocity vectors. Samples with speeds exceeding 5 standard deviations were classified as saccades. Blinks were detected automatically by the eye tracker. All remaining samples that were classified as neither blinks nor saccades were labeled as fixations. Because subsequent analyses focused exclusively on fixations that ended while the stimulus was still visible, the final fixation on each image was excluded.

#### MEG Data Acquisition and Preprocessing

To minimize head motion during recording, individualized head stabilizers were constructed for each participant. First, each participant’s head shape was captured using a Structure Sensor Pro (XRPro, LLC; https://structure.io) and compared with a 3D scan of the MEG helmet. The space between the participant’s head and the MEG headpiece was then milled into styrofoam to produce a personalized insert. Before each session, head shape was digitized using a Polhemus FASTRAK device (Polhemus; https://polhemus.com). To monitor head position within the MEG helmet, five head-position indicator coils were attached above the nasion on the forehead, bilaterally near the inions, and at intermediate forehead locations. Recordings were acquired in a magnetically shielded room using a 306-channel NeuroMag VectorView MEG system (Megin; https://megin.com), comprising 102 magnetometers and 204 planar gradiometers. Data were sampled at 1,000 Hz with an online low-pass filter at 330 Hz. Offline, raw MEG data were band-pass filtered between 0.2 and 200 Hz using a causal filter. Channels identified during acquisition as excessively noisy were monitored and interpolated before further analysis. Data were then denoised using spatiotemporal MaxFilter (Maxwell filtering; Elekta), including temporal Signal Space Separation (tSSS;[91]), which also compensated for differences in head position and movement across recordings. Subsequent analyses were performed in Python using MNE-Python [92] together with custom scripts. Independent component analysis (ICA) was applied separately to each participant and session to identify and remove noisy components. Components related to eye movements were identified by correlating fixation-locked component activity with the corresponding horizontal and vertical eye-position traces. To correct for session-specific offsets, MEG signals were median-centered separately for each channel and session.

#### Source reconstruction

Continuous MEG data were projected to source space using a linearly constrained minimum variance (LCMV) beamformer [93] implemented in MNE-Python [92]. Individual structural MRIs (3T) were processed with FreeSurfer [94] to reconstruct cortical surfaces and generate subject-specific head models. The forward solution was computed using a three-layer boundary element model (BEM), with the source space defined on the cortical surface using ico4 decimation (4098 vertices per hemisphere). Head stabilization was achieved using individually fitted foam casts during data collection. For each session, noise covariance was estimated from empty-room recordings acquired during each experimental session. Data covariance was computed empirically from the continuous raw recordings of each session. LCMV spatial filters were computed separately for each session using the corresponding data and noise covariance matrices (regularization = 0.05), with fixed source orientation constrained to the cortical surface normal, unit-noise-gain normalization, and rank determined from the data. Source activity was then reconstructed from the continuous data, yielding one signed source time course per vertex. Beta time courses were extracted using MNE’s extract label time course function. The ventral stream visual cortex, Medial temporal cortex, and Lateral temporal cortex regions were derived from the Glasser et al. (2016) anatomical parcellation [95]. Early visual cortex is used as defined in NSD [24], which encompasses V1, V2, and V3 from the Wang et al. (2015) [96] anatomical atlas.

#### Feature extraction

To assess which image features participants focused on at each fixation, fixation crops of 3.4° x 3.4° were extracted around each fixation participants made (Fig. 1a). From those crops a range of different image features were extracted, depending on the analysis.

##### SPoSE feature extraction

For the main analyses the dimpred model [32] was used to extract the SPoSE dimensions [28]. This model extracts a CLIP ResNet embedding [31] from each fixation crop, based on which it linearly predicts the SPoSE dimensions of each crop (Fig. 1b).

##### AlexNet feature extraction

For the AlexNet control analysis the fixation crops were passed through an AlexNet model [35] trained on ecoset [97]. We extracted activations from layers features.2 (low-level) and features.12 (high-level). For each layer, activations from all fixation crops across all subjects were concatenated and Principal Component Analysis (PCA) was applied. The first 25 PCs were retained for subsequent analyses.

##### Low-level image statistics

For the final control analysis, low-level visual properties were extracted from each fixation crop. Mean luminance was computed after converting each crop to grayscale using standard RGB luminance weights (0.299, 0.587, 0.114 for red, green, and blue channels, respectively), and averaging pixel intensities. Edge density was quantified by applying a Canny edge detector [98] to the grayscale crop and calculating the proportion of pixels classified as edges.

### Decoding analyses

For the decoding analysis MEG magnetometer data were epoched around each fixation onset (−200 to 500 ms). Within each experimental session, sensor data were z-scored separately for each sensor and time point across events, and data from all sessions were then concatenated. Each fixation event was matched to the SPoSE dimensions corresponding to its fixation crop and SPoSE dimensions were z-scored across samples. Decoding was performed separately for each dimension and each time point using ridge regression. At each time point the multivariate pattern across all MEG magnetometer sensors served as input, and one SPoSE dimension as the target variable. To prevent information leakage, data were split into training and validation sets at the level of scene identity, such that all samples from a given scene were assigned exclusively to one set. Model performance was quantified on the held-out validation set (20% of the data) using the Pearson correlation between predicted and observed dimension values, yielding time-resolved decoding accuracy for each dimension. To characterize similarities in temporal decoding dynamics across dimensions, time-resolved decoding accuracy curves were clustered using dynamic time warping (DTW). For each dimension, decoding time courses were first averaged across subjects. Dimensions with negligible decoding performance were excluded by applying a threshold on the maximum correlation (*≥* 0.05). To focus on temporal profile shape rather than absolute magnitude, time courses were z-scored per subject and dimension prior to averaging. Additionally, time courses were smoothed using a causal moving average filter (window length = 20 ms), where each time point was averaged over the current and preceding samples only, prior to DTW computation. Pairwise distances between dimension-specific time courses were computed using DTW, and hierarchical clustering was performed on the resulting distance matrix using complete linkage, following Teichmann et al. [30].

### Unfold analysis

Across all 50 sessions, linear deconvolution was applied to continuous MEG data, sampled at 500 Hz using Unfold [34]. The MEG data were z-scored across all sessions and sensors. For the main analyses fixation-onset was defined as the event. A supplementary analysis aligned to saccade onset is shown in Supplementary Fig. 5. The 49 z-scored SPoSE dimensions were included as continuous regressors, as well as fixation location, saccade amplitude, and saccade velocity to control for oculomotor artifacts. The same Unfold model was fit independently to each sensor (102 magnetometers) and, in a separate analysis, to each vertex of the source-reconstructed data (8196 vertices). Prior to model fitting, artifact-contaminated segments were identified using an adaptive peak-to-peak method: the maximum cross-channel peak-to-peak amplitude was computed in sliding 2-second windows (0.5 s step), and windows exceeding a data-driven threshold (median + 5 × MAD of the peak-to-peak distribution) were flagged as artifactual. Following [34] and [99], the corresponding rows of the time-expanded design matrix were set to zero rather than removed, preserving the temporal structure required for deconvolution. This procedure excluded on average 2.98% of the continuous data per session. Each session included on average 3936 fixation events. Fixation timing exhibited substantial overlap and variability, with median inter-event intervals of 310 ms across sessions relative to a modeled response window of −300 to 800 ms, indicating that overlapping responses were common and suitable for deconvolution. At the same time, inter-event intervals varied broadly, providing the temporal variability required for reliable separation of overlapping activity. Collinearity among the 49 SPoSE dimensions used as regressors was consistently moderate, with condition numbers of the z-scored predictor matrix averaging 4.99 and a mean maximum pairwise correlations of 0.6 across all sessions. Together, these diagnostics indicate that the design matrices were well-conditioned and that the single-session beta estimates were stably identifiable. Electrophysiological fields are typically bipolar over the magnetometer sensor array [100, 101] and the sign of source-localized MEG data is arbitrary due to dipole orientation ambiguity [102–104], which is why we did not interpret the polarity of the beta estimates. Furthermore, SPoSE dimensions capture diverse visual features that may modulate neural activity with different polarities which means that averaging beta estimates across dimensions risks cancellation. To prevent this, beta maps were aligned before averaging by calculating Pearson’s correlation coefficient between each dimension’s beta map and the beta map of the model intercept and multiplying the dimension’s map by −1 if the correlation was negative. The intercept captures the dominant spatial pattern of fixation-evoked activity, making it a natural reference for orienting dimension-specific modulations. The same approach was applied when averaging together multiple sensors from opposite sides of the head (e.g. Fig. 2d), which showed inverted signs due to the bipolar nature of magnetometer sensors. This approach has direct precedent in the MEG literature [103–105].

### Statistical testing

After running Unfold, beta estimates were analyzed using MNE [92] and custom Python scripts. All beta time courses were low-pass filtered at 50 Hz using a fourth-order Butterworth IIR filter applied in a causal manner, such that each time point depended only on current and past samples. The dataset comprised five independent participants (n = 5), each contributing ten recording sessions (50 sessions in total). Statistical analyses were performed on session-level beta estimates, with participant identity included as a random intercept to account for repeated measurements within participants. Statistical significance of beta estimates was assessed using linear mixedeffects models. At each time point and sensor, we fitted a model of the form estimate 1 + (1 — subject), where observations corresponded to session-level estimates. Models were fitted using maximum likelihood. Resulting p-values were corrected using the Benjamini–Hochberg false discovery rate (FDR) procedure [106], with a significance threshold of FDR-adjusted *p <* 0.05. For sensor-space analyses, FDR correction was performed across sensors and time bins. For source-space region-of-interest analyses, FDR correction was performed across all time points.

### GPN analysis

To quantify predictive uncertainty of visual input at a new location given a preceding sequence of saccades and fixations, we used the Glimpse Prediction Network [54]. This recurrent model is trained through self-supervision to predict the feature embedding of the next fixation patch from previous patches on the same image and relative saccade displacements (GPN variant GPN RS was used). First, fixation patches were passed through a ResNet50 [107] trained with the SimCLR objective [108], and activations were extracted after the average pool layer (n dimensions = 2,048). The ResNet50 embeddings of each fixation patch were then passed to GPN sequentially, according to the fixation sequence that the participants made on the image, along with the corresponding saccade vector. At each time step (i.e., each fixation), GPN predicted the embedding of the upcoming fixation patch, given the entire preceding fixation/sac-cade sequence. GPN activations were extracted at each time step at the later layer ‘lstm out’ (n dimensions = 2,048). For our analysis, we then quantified prediction error as the cosine distance between the GPN’s predicted embedding and the true ResNet50 embedding of a particular fixation patch.

## Data availability

The MEG, eye-tracking, and structural MRI data analyzed in this study are publicly available through the Active Visual Semantics (AVS) dataset [27] under a Creative Commons Attribution 4.0 International (CC BY 4.0) licence at https://registry.opendata.aws/avs/. The dataset comprises ten recording sessions for each of five participants and is accessible via the S3 bucket kietzmannlab-avs.

## Code availability

Custom code used for feature extraction, MEG decoding, linear deconvolution, source-level analyses, statistical testing and figure generation is available at https://github.com/Lucakaemmer/PredictiveNaturalVision.

## Acknowledgements

We thank Adrien Doerig, Sushrut Thorat, and Sander van Bree for their insightful feedback. This work was supported by the ERC Starting Grant TIME (grant no. 101039524 to T.C.K.), the Studienstiftung des Deutschen Volkes (to L.K.), a Max Planck Society research group grant (to M.N.H.), the ERC Starting Grant COREDIM (grant no. ERC-2021-STG-101039712 to M.N.H.), a LOEWE Start Professorship from the Hessian Ministry of Higher Education, Research, Science and the Arts (to M.N.H.) and the Deutsche Forschungsgemeinschaft under Germany’s Excellence Strategy (EXC 3066/1 ‘The Adaptive Mind’, grant no. 533717223 to M.N.H.). The Osnabrück HPC received support from the Deutsche Forschungsgemeinschaft (grant no. 456666331).

## Author contributions

L.K. contributed to conceptualization, methodology, data curation, formal analysis, visualization, and writing the original draft. C.A. contributed to conceptualization, methodology, investigation, data curation, and writing – review and editing. P.S. contributed to conceptualization, methodology, investigation, data curation, and writing – review and editing. M.N.H. contributed to methodology, supervision, funding acquisition, and writing – review and editing. T.K. contributed to conceptualization, methodology, supervision, funding acquisition, resources, and writing – review and editing. All authors read and approved the final manuscript.

## References

[1] Summerfield, C., Trittschuh, E. H., Monti, J. M., Mesulam, M.-M. & Egner, T. Neural repetition suppression reflects fulfilled perceptual expectations. Nature neuroscience 11, 1004–1006 (2008).

[2] Kok, P., Jehee, J. F. & De Lange, F. P. Less is more: expectation sharpens representations in the primary visual cortex. Neuron 75, 265–270 (2012).

[3] Kok, P., Failing, M. F. & de Lange, F. P. Prior expectations evoke stimulus templates in the primary visual cortex. Journal of cognitive neuroscience 26, 1546–1554 (2014).

[4] De Lange, F. P., Heilbron, M. & Kok, P. How do expectations shape perception? Trends in cognitive sciences 22, 764–779 (2018).

[5] Todorovic, A. & Auksztulewicz, R. Dissociable neural effects of temporal expectations due to passage of time and contextual probability. Hearing research 399, 107871 (2021).

[6] Egner, T., Monti, J. M. & Summerfield, C. Expectation and surprise deter-mine neural population responses in the ventral visual stream. Journal of Neuroscience 30, 16601–16608 (2010).

[7] Kämmer, L., Kroell, L. M., Knapen, T., Rolfs, M. & Hebart, M. N. Feedback of peripheral saccade targets to early foveal cortex. eLife 14, RP107053 (2026).

[8] Summerfield, C. & Egner, T. Expectation (and attention) in visual cognition. Trends in cognitive sciences 13, 403–409 (2009).

[9] Stefanics, G., Kremláček, J. & Czigler, I. Visual mismatch negativity: a predictive coding view. Frontiers in human neuroscience 8, 666 (2014).

[10] Kok, P. & de Lange, F. P. Predictive coding in sensory cortex An introduction to model-based cognitive neuroscience (eds Forstmann, B. U. & Wagenmakers, E.-J.) 221–244 (Springer, 2015).

[11] Heilbron, M. & de Lange, F. P. Higher-level spatial prediction in natural vision across mouse visual cortex. PLOS Computational Biology 22, e1013136 (2026).

[12] Yarbus, A. L. Eye movements during perception of complex objects Eye movements and vision 171–211 (Springer, 1967).

[13] Findlay, J. M. & Gilchrist, I. D. Active vision: The psychology of looking and seeing 37 (Oxford University Press, 2003).

[14] Rolfs, M., Jonikaitis, D., Deubel, H. & Cavanagh, P. Predictive remapping of attention across eye movements. Nature neuroscience 14, 252–256 (2011).

[15] Schroeder, C. E., Wilson, D. A., Radman, T., Scharfman, H. & Lakatos, P. Dynamics of active sensing and perceptual selection. Current opinion in neurobiology 20, 172–176 (2010).

[16] Hayhoe, M. & Ballard, D. Eye movements in natural behavior. Trends in cognitive sciences 9, 188–194 (2005).

[17] Kietzmann, T. C. & König, P. Effects of contextual information and stimulus ambiguity on overt visual sampling behavior. Vision research 110, 76–86 (2015).

[18] Rayner, K. Eye movements in reading and information processing: 20 years of research. Psychological bulletin 124, 372–422 (1998).

[19] Torralba, A., Oliva, A., Castelhano, M. S. & Henderson, J. M. Contextual guidance of eye movements and attention in real-world scenes: the role of global features in object search. Psychological review 113, 766–786 (2006).

[20] Hwang, A. D., Wang, H.-C. & Pomplun, M. Semantic guidance of eye movements in real-world scenes. Vision research 51, 1192–1205 (2011).

[21] Kroner, A., Senden, M., Driessens, K. & Goebel, R. Contextual encoder–decoder network for visual saliency prediction. Neural Networks 129, 261–270 (2020).

[22] Henderson, J. M. & Hayes, T. R. Meaning-based guidance of attention in scenes as revealed by meaning maps. Nature human behaviour 1, 743–747 (2017).

[22] Kämmer, L., Kroner, A. & Hebart, M. N. Interpretable prediction of human fixations from behavior-derived representational dimensions. Poster presentation, 8th Annual Conference on Cognitive Computational Neuroscience (CCN) (2025). URL https://2025.ccneuro.org/poster/?id=ieHdINQtCb.

[24] Allen, E. J. et al. A massive 7t fmri dataset to bridge cognitive neuroscience and artificial intelligence. Nature neuroscience 25, 116–126 (2022).

[25] Hebart, M. N. et al. Things-data, a multimodal collection of large-scale datasets for investigating object representations in human brain and behavior. Elife 12, e82580 (2023).

[26] DiCarlo, J. J., Zoccolan, D. & Rust, N. C. How does the brain solve visual object recognition? Neuron 73, 415–434 (2012).

[27] Sulewski, P., Amme, C., König, P., Hebart, M. N. & Kietzmann, T. C. Active visual semantics: A large-scale meg and eye-tracking dataset for understanding visual intelligence in action. arXiv preprint arXiv:2609.01055 (2026).

[28] Hebart, M. N., Zheng, C. Y., Pereira, F. & Baker, C. I. Revealing the multidimensional mental representations of natural objects underlying human similarity judgements. Nature human behaviour 4, 1173–1185 (2020).

[29] Contier, O., Baker, C. I. & Hebart, M. N. Distributed representations of behaviour-derived object dimensions in the human visual system. Nature Human Behaviour 8, 2179–2193 (2024).

[30] Teichmann, L., Hebart, M. N. & Baker, C. I. Dynamic representation of multidimensional object properties in the human brain. Journal of Neuroscience 46, e1057252026 (2026).

[31] Radford, A. et al. Learning transferable visual models from natural language supervision. International conference on machine learning, 8748–8763 (PmLR, 2021).

[32] Kaniuth, P., Mahner, F. P., Perkuhn, J. & Hebart, M. N. A high-throughput approach for the efficient prediction of perceived similarity of natural objects. BioRxiv 2024–06 (2024).

[33] Kramer, M. A., Hebart, M. N., Baker, C. I. & Bainbridge, W. A. The features underlying the memorability of objects. Science advances 9, eadd2981 (2023).

[34] Ehinger, B. V. & Dimigen, O. Unfold: an integrated toolbox for overlap correction, non-linear modeling, and regression-based eeg analysis. PeerJ 7, e7838 (2019).

[35] Krizhevsky, A., Sutskever, I. & Hinton, G. E. Imagenet classification with deep convolutional neural networks. Advances in neural information processing systems 25 (2012).

[36] Zeiler, M. D. & Fergus, R. Visualizing and understanding convolutional net-works. European conference on computer vision, 818–833 (Springer, 2014).

[37] Yosinski, J., Clune, J., Nguyen, A., Fuchs, T. & Lipson, H. Understanding neural networks through deep visualization. arXiv preprint arXiv:1506.06579 (2015).

[38] Yamins, D. L. et al. Performance-optimized hierarchical models predict neural responses in higher visual cortex. Proceedings of the national academy of sciences 111, 8619–8624 (2014).

[39] Thorpe, S., Fize, D. & Marlot, C. Speed of processing in the human visual system. nature 381, 520–522 (1996).

[40] Di Russo, F., Martínez, A., Sereno, M. I., Pitzalis, S. & Hillyard, S. A. Cortical sources of the early components of the visual evoked potential. Human brain mapping 15, 95–111 (2002).

[41] Liu, J., Harris, A. & Kanwisher, N. Stages of processing in face perception: an MEG study. Nature Neuroscience 5, 910–916 (2002). URL 10.1038/nn909.

[42] Bar, M. The proactive brain: using analogies and associations to generate predictions. Trends in cognitive sciences 11, 280–289 (2007).

[43] Kok, P., Mostert, P. & De Lange, F. P. Prior expectations induce prestimulus sensory templates. Proceedings of the National Academy of Sciences 114, 10473– 10478 (2017).

[44] Hindy, N. C., Ng, F. Y. & Turk-Browne, N. B. Linking pattern completion in the hippocampus to predictive coding in visual cortex. Nature neuroscience 19, 665–667 (2016).

[45] Kok, P. & Turk-Browne, N. B. Associative prediction of visual shape in the hippocampus. Journal of Neuroscience 38, 6888–6899 (2018).

[46] Kutas, M. & Hillyard, S. A. Reading senseless sentences: Brain potentials reflect semantic incongruity. Science 207, 203–205 (1980).

[47] Kutas, M. & Hillyard, S. A. Brain potentials during reading reflect word expectancy and semantic association. Nature 307, 161–163 (1984).

[48] Hamm, J. P., Johnson, B. W. & Kirk, I. J. Comparison of the n300 and n400 erps to picture stimuli in congruent and incongruent contexts. Clinical neurophysiology 113, 1339–1350 (2002).

[49] Cohn, N., Paczynski, M., Jackendoff, R., Holcomb, P. J. & Kuperberg, G. R. (pea) nuts and bolts of visual narrative: Structure and meaning in sequential image comprehension. Cognitive psychology 65, 1–38 (2012).

[50] Osterhout, L. & Holcomb, P. J. Event-related brain potentials elicited by syntactic anomaly. Journal of memory and language 31, 785–806 (1992).

[51] Sitnikova, T., Holcomb, P. J. & Kuperberg, G. R. Neurocognitive mechanisms of human comprehension. Understanding events: How humans see, represent, and act on events 639–683 (2008).

[52] Mudrik, L., Lamy, D. & Deouell, L. Y. Erp evidence for context congruity effects during simultaneous object–scene processing. Neuropsychologia 48, 507–517 (2010).

[53] Vo, M. L.-H. & Wolfe, J. M. Differential electrophysiological signatures of semantic and syntactic scene processing. Psychological science 24, 1816–1823 (2013).

[54] Thorat, S., Doerig, A., Kroner, A., Amme, C. & Kietzmann, T. C. Predicting upcoming visual features during eye movements yields scene representations aligned with human visual cortex. arXiv:2511.12715 (2025). URL https://arxiv.org/abs/2511.12715. Version Number: 1.

[55] Sperry, R. W. Neural basis of the spontaneous optokinetic response produced by visual inversion. Journal of comparative and physiological psychology 43, 482–489 (1950).

[56] Sommer, M. A. & Wurtz, R. H. A pathway in primate brain for internal monitoring of movements. science 296, 1480–1482 (2002).

[57] Sommer, M. A. & Wurtz, R. H. What the brain stem tells the frontal cortex. ii. role of the sc-md-fef pathway in corollary discharge. Journal of neurophysiology 91, 1403–1423 (2004).

[58] Grill-Spector, K., Henson, R. & Martin, A. Repetition and the brain: neural models of stimulus-specific effects. Trends in cognitive sciences 10, 14–23 (2006).

[59] Kimura, M., Schröger, E. & Czigler, I. Visual mismatch negativity and its importance in visual cognitive sciences. Neuroreport 22, 669–673 (2011).

[60] Carmel R, A.-A., Gal, V., Oded, W. & Leon Y, D. Decoding object categories from eeg during free viewing reveals early information evolution compared to passive viewing. BioRxiv 2023–06 (2023).

[61] Fakche, C., Hickey, C. & Jensen, O. Fast feature-and category-related parafoveal previewing support free visual exploration. Journal of Neuroscience 44, e0841242024 (2024).

[62] Buonocore, A., Dimigen, O. & Melcher, D. Post-saccadic face processing is modulated by pre-saccadic preview: Evidence from fixation-related potentials. Journal of Neuroscience 40, 2305–2313 (2020).

[63] Herwig, A. & Schneider, W. X. Predicting object features across saccades: evidence from object recognition and visual search. Journal of Experimental Psychology: General 143, 1903–1922 (2014).

[64] Valsecchi, M. & Gegenfurtner, K. R. Dynamic re-calibration of perceived size in fovea and periphery through predictable size changes. Current Biology 26, 59–63 (2016).

[65] Cichy, R. M., Pantazis, D. & Oliva, A. Resolving human object recognition in space and time. Nature neuroscience 17, 455–462 (2014).

[66] Dumoulin, S. O. & Wandell, B. A. Population receptive field estimates in human visual cortex. Neuroimage 39, 647–660 (2008).

[67] Gert, A. L., Ehinger, B. V., Timm, S., Kietzmann, T. C. & König, P. Wildlab: A naturalistic free viewing experiment reveals previously unknown electroencephalography signatures of face processing. European Journal of Neuroscience 56, 6022–6038 (2022).

[68] de Vries, I. E. & Wurm, M. F. Predictive neural representations of naturalistic dynamic input. Nature Communications 14, 3858 (2023).

[69] Richter, D., Kietzmann, T. C. & de Lange, F. P. High-level visual prediction errors in early visual cortex. PLoS biology 22, e3002829 (2024).

[70] Zhang, M., et al. From thought to action: How a hierarchy of neural dynamics supports language production. arXiv preprint arXiv:2502.07429 (2025).

[71] Kveraga, K., Ghuman, A. S. & Bar, M. Top-down predictions in the cognitive brain. Brain and cognition 65, 145–168 (2007).

[72] Deubel, H. & Schneider, W. X. Saccade target selection and object recognition: Evidence for a common attentional mechanism. Vision research 36, 1827–1837 (1996).

[73] Kowler, E., Anderson, E., Dosher, B. & Blaser, E. The role of attention in the programming of saccades. Vision research 35, 1897–1916 (1995).

[74] Li, H.-H., Barbot, A. & Carrasco, M. Saccade preparation reshapes sensory tuning. Current Biology 26, 1564–1570 (2016).

[75] Wang, Y., Cao, R. & Wang, S. Encoding of visual objects in the human medial temporal lobe. Journal of Neuroscience 44, e2135232024 (2024).

[76] Rolls, E. T. et al. A ventromedial visual cortical ‘where’stream to the human hippocampus for spatial scenes revealed with magnetoencephalography. Communications Biology 7, 1047 (2024).

[77] Huth, A. G., De Heer, W. A., Griffiths, T. L., Theunissen, F. E. & Gallant, J. L. Natural speech reveals the semantic maps that tile human cerebral cortex. Nature 532, 453–458 (2016).

[78] Mickiewicz, E. A. et al. A semantotopic map in human hippocampus. bioRxiv 2025–10 (2025).

[79] Clarke, A. Neural dynamics of visual and semantic object processing Psychology of learning and motivation, Vol. 70 71–95 (Elsevier, 2019).

[80] Clarke, A. & Tyler, L. K. Object-specific semantic coding in human perirhinal cortex. The Journal of neuroscience 34, 4766–4775 (2014).

[81] Clarke, A., Devereux, B. J. & Tyler, L. K. Oscillatory dynamics of perceptual to conceptual transformations in the ventral visual pathway. Journal of cognitive neuroscience 30, 1590–1605 (2018).

[82] Bonnen, T., Yamins, D. L. & Wagner, A. D. When the ventral visual stream is not enough: A deep learning account of medial temporal lobe involvement in perception. Neuron 109, 2755–2766 (2021).

[83] Warrington, O., Graedel, N. N., Callaghan, M. F. & Kok, P. Communication of perceptual predictions from the hippocampus to the deep layers of the parahippocampal cortex. Science Advances 11, eads4970 (2025).

[84] Raghavan, V. S. et al. During natural vision, semantic novelty modulates fixation-related processing in primate cortex. bioRxiv 2026–03 (2026).

[85] Pizzo, F. et al. Deep brain activities can be detected with magnetoencephalography. Nature communications 10, 971 (2019).

[86] Ruzich, E., Crespo-García, M., Dalal, S. S. & Schneiderman, J. F. Characterizing hippocampal dynamics with meg: A systematic review and evidence-based guidelines. Human brain mapping 40, 1353–1375 (2019).

[87] Stolk, A., Todorovic, A., Schoffelen, J.-M. & Oostenveld, R. Online and offline tools for head movement compensation in meg. Neuroimage 68, 39–48 (2013).

[88] Zerbe, J., et al. LAION-fMRI: A densely sampled 7t-fmri dataset providing broad coverage of natural image diversity (2026). URL https://www.visionsciences.org/talk-sessions?id=642. Talk 25.11, Scene Perception Talk Session, Vision Sciences Society Annual Meeting, May 16, 2026.

[89] Lin, T.-Y. et al. Microsoft COCO: Common objects in context. Computer Vision – ECCV 2014 (eds Fleet, D., Pajdla, T., Schiele, B. & Tuytelaars, T.), 740–755 (Springer, 2014). URL 10.1007/978-3-319-10602-148.

[90] Engbert, R. & Mergenthaler, K. Microsaccades are triggered by low retinal image slip. Proceedings of the National Academy of Sciences 103, 7192–7197 (2006). URL https://www.pnas.org/doi/10.1073/pnas.0509557103.

[91] Taulu, S. & Simola, J. Spatiotemporal signal space separation method for rejecting nearby interference in MEG measurements. Physics in medicine and biology 51, 1759–1768 (2006). URL https://pubmed.ncbi.nlm.nih.gov/16552102/.

[92] Gramfort, A. et al. MEG and EEG data analysis with MNE-python. Frontiers in Neuroscience 7, 267 (2013).

[93] Van Veen, B., Van Drongelen, W., Yuchtman, M. & Suzuki, A. Localization of brain electrical activity via linearly constrained minimum variance spatial filtering. IEEE Transactions on Biomedical Engineering 44, 867–880 (1997). URL http://ieeexplore.ieee.org/document/623056/.

[94] Fischl, B. FreeSurfer. NeuroImage 62, 774–781 (2012). URL https://www.ncbi.nlm.nih.gov/pmc/articles/PMC3685476/.

[95] Glasser, M. F. et al. A multi-modal parcellation of human cerebral cortex. Nature 536, 171–178 (2016).

[96] Wang, L., Mruczek, R. E., Arcaro, M. J. & Kastner, S. Probabilistic maps of visual topography in human cortex. Cerebral cortex 25, 3911–3931 (2015).

[97] Mehrer, J., Spoerer, C. J., Jones, E. C., Kriegeskorte, N. & Kietzmann, T. C. An ecologically motivated image dataset for deep learning yields better models of human vision. Proceedings of the National Academy of Sciences 118, e2011417118 (2021). URL https://pnas.org/doi/full/10.1073/pnas.2011417118.

[98] Canny, J. A computational approach to edge detection. IEEE Transactions on Pattern Analysis and Machine Intelligence PAMI-8, 679–698 (1986). URL 10.1109/TPAMI.1986.4767851.

[99] Smith, N. J. & Kutas, M. Regression-based estimation of erp waveforms: Ii. non-linear effects, overlap correction, and practical considerations. Psychophysiology 52, 169–181 (2015).

[100] Hämäläinen, M., Hari, R., Ilmoniemi, R. J., Knuutila, J. & Lounasmaa, O. V. Magnetoencephalography—theory, instrumentation, and applications to noninvasive studies of the working human brain. Reviews of modern Physics 65, 413–497 (1993).

[101] Gross, J. et al. Good practice for conducting and reporting meg research. Neuroimage 65, 349–363 (2013).

[102] Colclough, G. L., Brookes, M. J., Smith, S. M. & Woolrich, M. W. A symmetric multivariate leakage correction for meg connectomes. Neuroimage 117, 439–448 (2015).

[103] Takagi, Y., Hunt, L. T., Woolrich, M. W., Behrens, T. E. & Klein-Flügge, M. C. Adapting non-invasive human recordings along multiple task-axes shows unfolding of spontaneous and over-trained choice. Elife 10, e60988 (2021).

[104] Brookes, M. J. et al. A multi-layer network approach to meg connectivity analysis. Neuroimage 132, 425–438 (2016).

[105] Huber, D. E., Tian, X., Curran, T., O’Reilly, R. C. & Woroch, B. The dynamics of integration and separation: Erp, meg, and neural network studies of immediate repetition effects. Journal of Experimental Psychology: Human Perception and Performance 34, 1389–1416 (2008).

[106] Benjamini, Y. & Hochberg, Y. Controlling the false discovery rate: a practical and powerful approach to multiple testing. Journal of the Royal Statistical Society: Series B (Methodological) 57, 289–300 (1995). URL 10.1111/j.2517-6161.1995.tb02031.x.

[107] He, K., Zhang, X., Ren, S. & Sun, J. Deep residual learning for image recognition. 2016 IEEE Conference on Computer Vision and Pattern Recognition (CVPR), 770–778 (IEEE, Las Vegas, NV, USA, 2016). URL http://ieeexplore.ieee.org/document/7780459/.

[108] Chen, T., Kornblith, S., Norouzi, M. & Hinton, G. A simple framework for contrastive learning of visual representations. Proceedings of the 37th Inter-national Conference on Machine Learning, 1597–1607 (PMLR, 2020). URL https://proceedings.mlr.press/v119/chen20j.html.

