## Supplementary Information for "Humans predictively process high-level information during active natural vision"

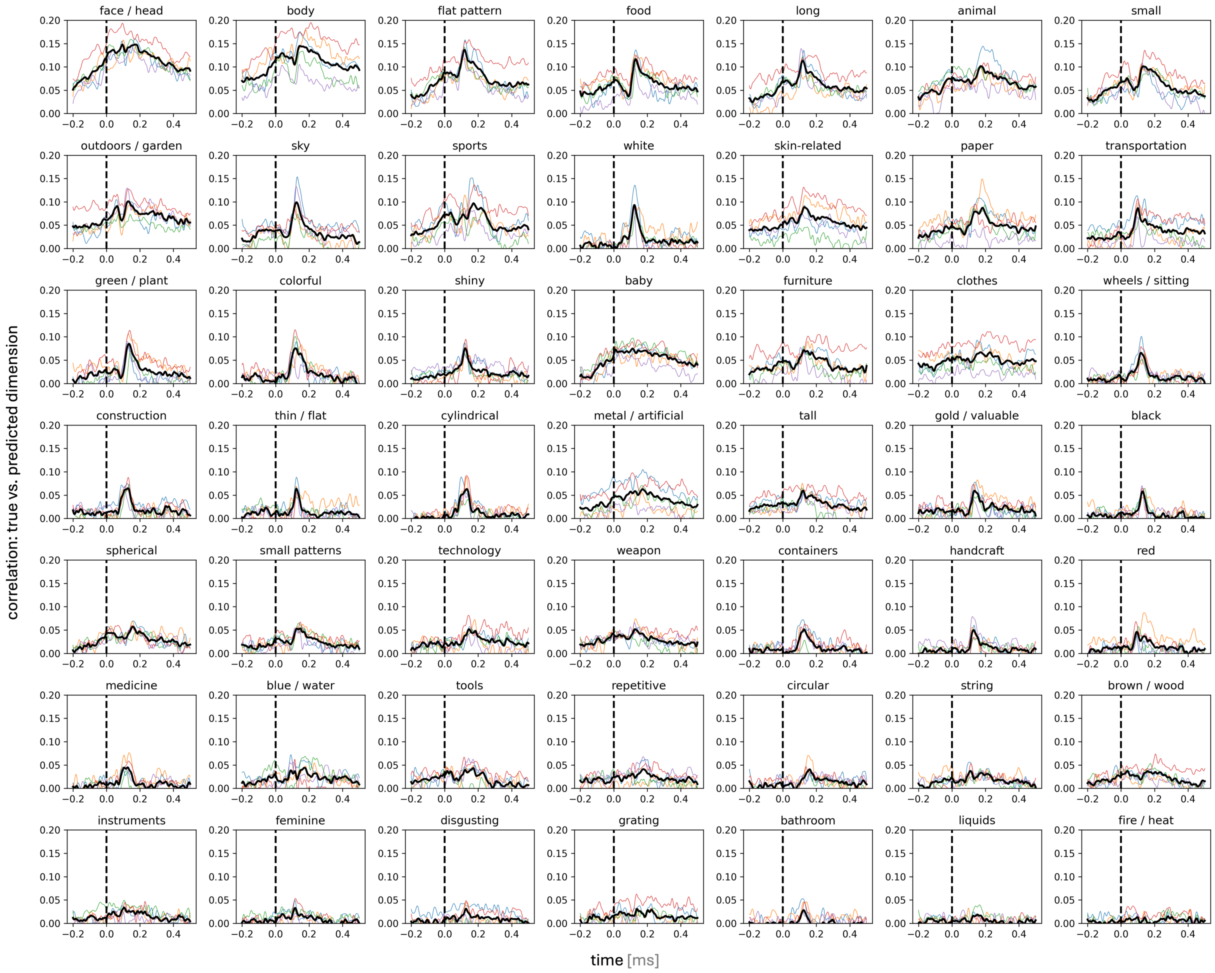


**Supplementary Fig. 1. Decoding time courses of all SPoSE dimensions.** Dimensions are sorted by peak decoding accuracy.

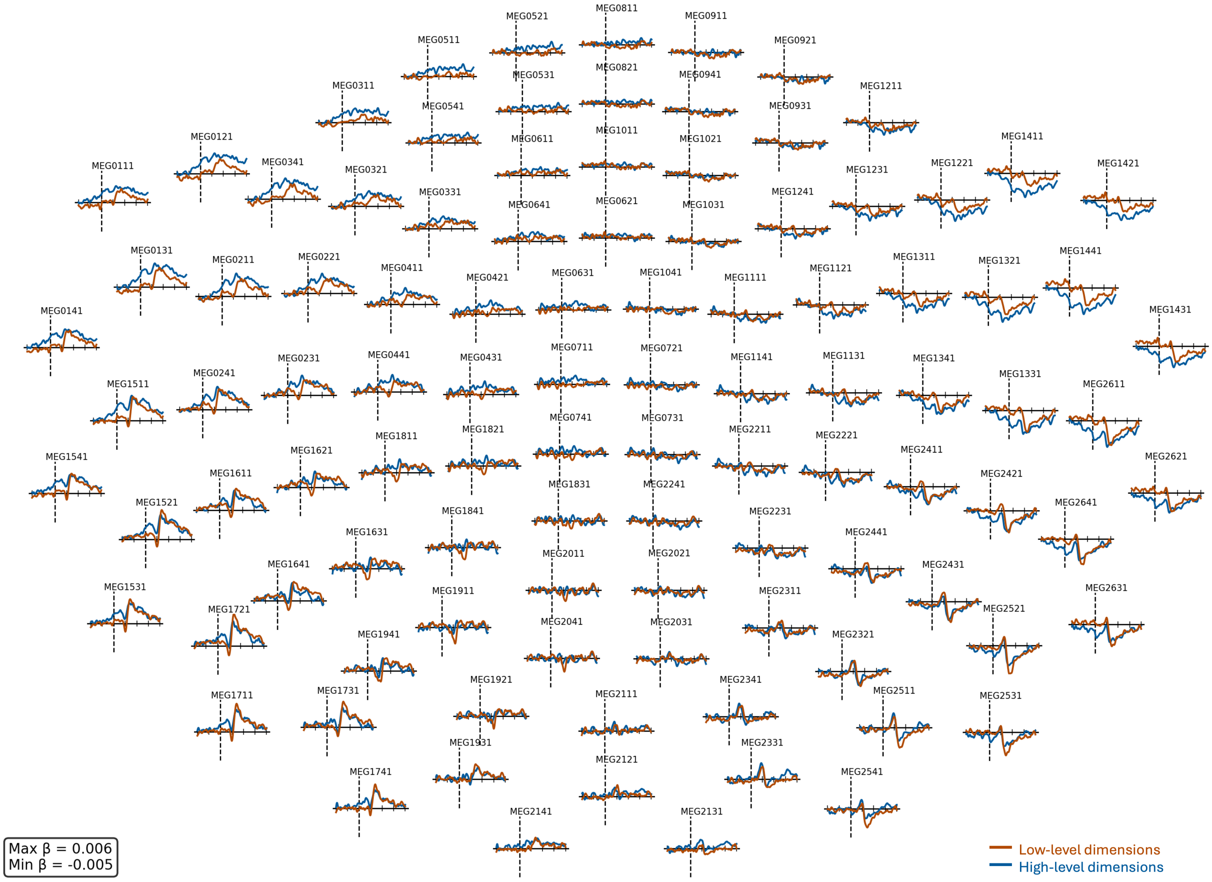


**Supplementary Fig. 2. Unfold beta estimates across MEG sensors.** Beta estimates are shown for all magnetometer sensors, averaged separately across low- and high-level SPoSE dimensions.

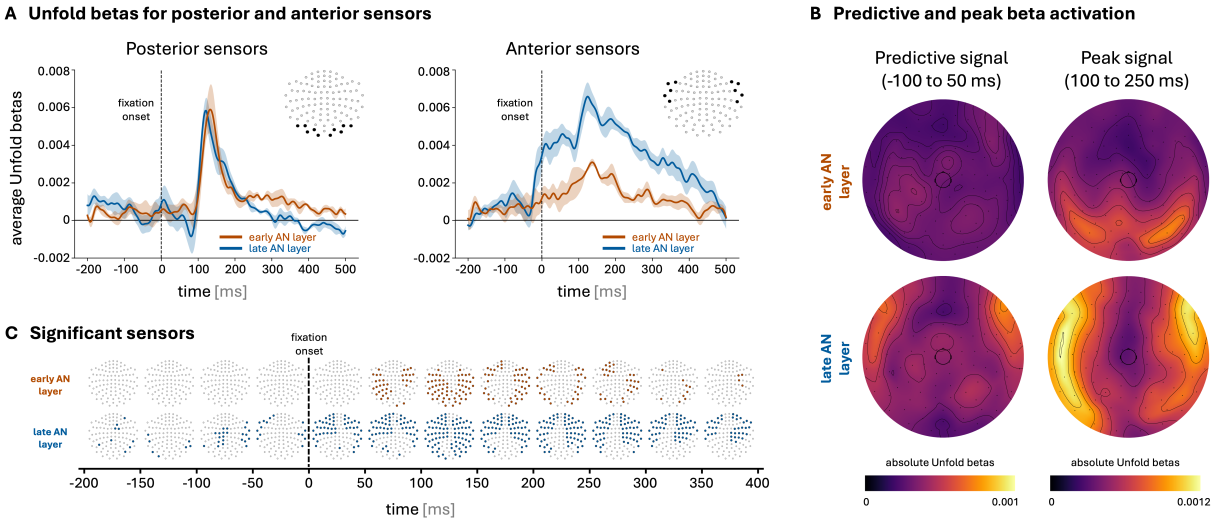


**Supplementary Fig. 3. Temporal modulation of low- and high-level AlexNet features.** The main Unfold sensor analysis was repeated using 25 principal components extracted from early and late AlexNet layers, respectively. All other analysis steps were identical to the SPoSE-based sensor analysis (Fig. 2 in the main manuscript).

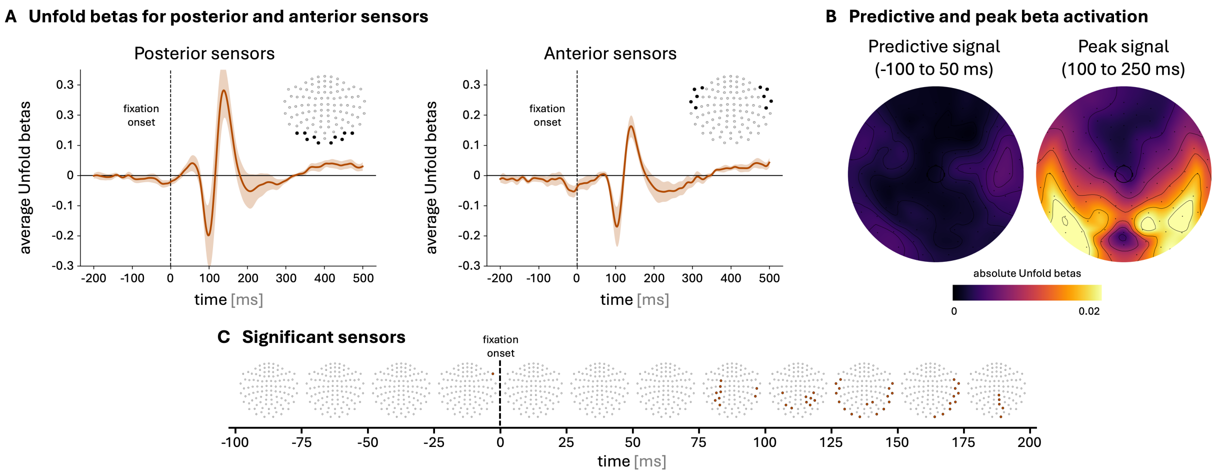


**Supplementary Fig. 4. Temporal modulation of low-level image properties.** The main Unfold sensor analysis was repeated using luminance and edge density extracted from fixation crops. All other analysis steps were identical to the SPoSE-based sensor analysis (Fig. 2 in the main manuscript).

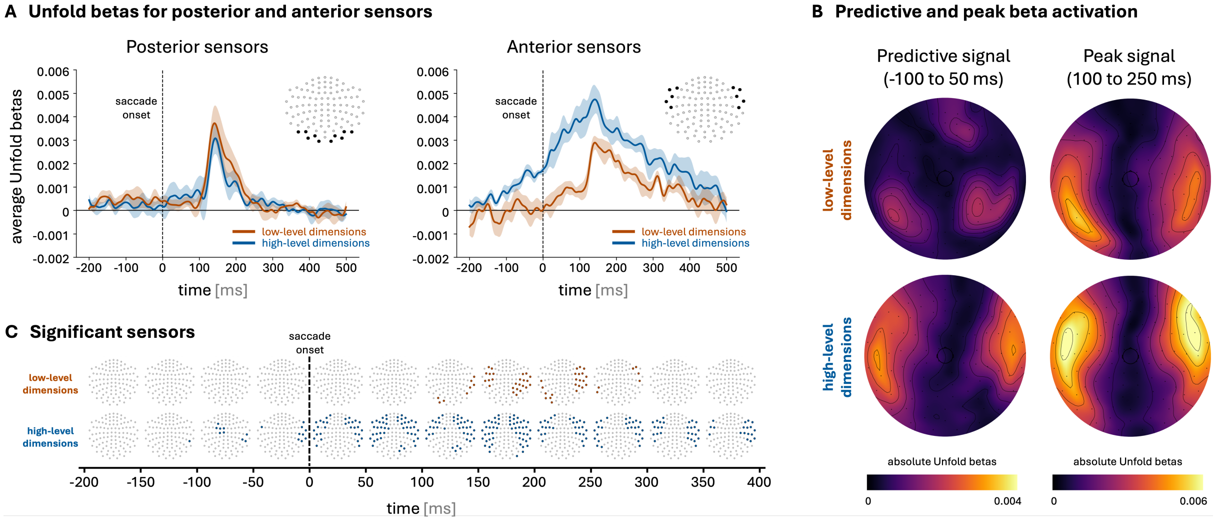


**Supplementary Fig. 5. Saccade-onset-aligned Unfold analysis.** The main Unfold sensor analysis was repeated using saccade onset instead of fixation onset as the event. All other analysis steps were identical to the fixation-onset-aligned SPoSE analysis (Fig. 2 in the main manuscript).
